# Defining critical drivers of cross-pollination for better hybrid grain set in wheat

**DOI:** 10.64898/2026.06.01.729316

**Authors:** Deepak Kumar, Monika Spiller, Laurent Gouere, Thorsten Schnurbusch

**Author notes:** corresponding authors: Deepak Kumar, Thorsten Schnurbusch.

## Abstract

Hybrid wheat breeding offers a promising route to enhance grain yield and yield stability through heterosis, yet hybrid grain production remains constrained by limited cross-pollination efficiency due to high rates of autogamy. To achieve cross-pollination in an autogamous species like wheat, pollen must shed outside the floret. This is typically assessed by scoring visual anther extrusion (VAEX), a key floral trait that sets the foundation for cross-pollination. However, VAEX explains only part of the variation in hybrid grain set. To address this, we analyzed floral structures and reproductive processes underlying cross-pollination efficiency in wheat. From 24 elite winter wheat genotypes, we developed traits describing anther extrusion kinetics, pollen release, and floral bract architecture. These traits showed substantial genotypic variation and high heritability. While VAEX alone explained approximately 49% of the variation in hybrid grain set, combined trait analyses explained up to 77%, demonstrating that hybrid grain production is governed by coordinated floral and reproductive trait interactions. Together, our analyses define a hierarchical trait architecture linking floral bract mechanics, anther extrusion dynamics, and pollen shedding to cross-fertilization success. This establishes a systems-level phenotyping framework for improving male parent selection in hybrid wheat breeding.

**Highlight:** High cross-pollination efficiency in wheat is a multi-factorial process that requires lighter floral bract architecture combined with adequate anther extrusion and pollen release for improving hybrid grain production.

## Introduction

Global wheat improvement for grain yield is approaching a biological and technological inflection point. After decades of steady progress driven by conventional breeding, yield gains in bread wheat (*Triticum aestivum* L.) have slowed despite continued genetic and agronomic innovation, exposing limitations of selection strategies based primarily on recombination within increasingly narrow germplasm pools (Evenson and Gollin, 2003; Ray *et al*., 2013; Tadesse *et al*., 2019). At the same time, increasing climatic instability and growing food demand are widening the gap between achievable and required wheat productivity, prompting renewed interest in breeding strategies capable of delivering step changes in grain yield rather than incremental improvements (Basnet *et al*., 2022; Shiferaw *et al*., 2013).

Hybrid breeding represents one such strategy. By exploiting heterosis, hybrids have transformed productivity in cereals such as maize and rice, producing substantial gains in yield potential, resilience, and yield stability across environments (Duvick, 2005; Labroo *et al*., 2021; Virmani, 2003). Wheat, however, has been more difficult to establish as a widely adopted hybrid crop (Rohde *et al*., 2025). Although fertility-control systems, genomic tools for parent selection, and hybrid breeding pipelines have advanced considerably, the realized grain yield advantages of wheat hybrids have generally remained more modest than those reported for several other cereals (Basnet *et al*., 2022; Longin *et al*., 2013; Whitford *et al*., 2013). Beyond achieving sufficient heterosis, the broader deployment of hybrid wheat also depends on producing hybrids efficiently and economically at scale. This requirement remains a major constraint, as hybrid production efficiency is strongly influenced by the cross-pollination capacity of parental lines. Consequently, limited male outcrossing ability currently restricts the choice of parents and reduces the broader exploitation of heterosis in wheat. This significant limitation is closely linked to the reproductive biology of wheat. Wheat is predominantly autogamous, with florets that frequently self-pollinate before opening, limiting opportunities for cross-fertilization (Selva *et al*., 2020; Whitford *et al*., 2013). Cleistogamy, i.e. limited floret opening, and imperfect flowering synchrony further restrict pollen transfer (Schmidt *et al*., 2025; Ueno and Itoh, 1997). In addition, wheat produces trinucleate pollen that is highly sensitive to desiccation and exhibits short longevity under field conditions, reducing the effective window for fertilization (Heslop-Harrison, 1968; Impe *et al*., 2020; Pacini and Dolferus, 2019). Consequently, hybrid grain production in wheat depends critically on efficient pollen release and dispersal during anthesis (Rohde *et al*., 2025).

Anthesis in wheat requires a fine-tuned orchestration among multiple floral organs; e.g. lodicule swelling drives floret opening, filament elongation positions anthers, and anther dehiscence enables pollen release (Browne *et al*., 2018; Selva *et al*., 2020). The degree to which anthers emerge beyond the enclosing lemma and palea determines pollen exposure to air currents and thus cross-pollination efficiency (Zajączkowska *et al*., 2021). Among floral traits, anther extrusion, scored as visual anther extrusion (VAEX) during anthesis, sets the basic anatomical precondition for any cross-pollination; and therefore, VAEX has become so far the main phenotypic criterion used to evaluate male parental lines in hybrid breeding programs (Boeven *et al*., 2016; El Hanafi *et al*., 2022). VAEX is typically scored as the proportion of extruded anthers visible outside the floret and provides a rapid field-based estimate of pollen dispersal potential.

The adoption of VAEX substantially improved male parent selection by linking male floral morphology to hybrid grain production (Boeven *et al*., 2016; El Hanafi *et al*., 2022). However, important limitations remain. VAEX phenotyping is labor-intensive, environmentally sensitive, and prone to observer bias, resulting in substantial experimental variability (Adhikari *et al*., 2020; Selva *et al*., 2020; Winn *et al*., 2021). Moreover, VAEX explains only a moderate proportion of variation in hybrid grain set, indicating that extrusion magnitude alone does not fully capture cross-pollination efficiency (Muqaddasi *et al*., 2017). This limitation becomes evident when genotypes with contrasting VAEX phenotypes produce comparable amounts of hybrid grains, suggesting that additional floral traits contribute to cross-pollination success (Boeven *et al*., 2016; Muqaddasi *et al*., 2017; Selva *et al*., 2020).

Despite the central role of floral biology in hybrid grain production, critical aspects of wheat anthesis remain insufficiently defined (Rohde *et al*., 2025). Previous studies have focused primarily on pollen viability and pollen number (Impe *et al*., 2020; Kanwal *et al*., 2022), whereas traits describing the dynamics and mechanics of pollen export have received comparatively little attention. Successful fertilization depends not only on pollen production but also on efficient pollen release, extrusion dynamics, and the structural properties of floral organs that regulate floret opening. Identifying and integrating such traits into wheat breeding pipelines represents a major opportunity to improve hybrid grain set and refine male parental line characterization (Rohde *et al*., 2025).

To address this, we investigated whether quantitative characterization of floral architecture and pollen dispersal dynamics improve hybrid grain set in wheat beyond VAEX. By exerting detailed phenotyping of 24 elite winter wheat genotypes combined with multivariate statistical modeling, we dissected the trait relationships among anther extrusion kinetics, pollen dispersal dynamics, and floral bract traits during anthesis. Our analyses reveal that hybrid grain production in wheat is determined by a series of hierarchical trait determinants of cross-pollination success rather than a single morphological indicator, with distinct traits contributing through both direct and indirect pathways. By quantifying these contributions, we establish a multi-factorial framework for elucidating hybrid grain set and refine the criteria used to characterize male parental lines in hybrid wheat breeding. Together, our findings shift phenotyping for hybrid wheat from single-trait evaluation toward a systems-level understanding of floral reproductive function, providing a mechanistic foundation for improved hybrid wheat breeding.

## Materials and Methods

### Plant material and experimental design

In total 24 elite bread wheats (*Triticum aestivum* L.), intended for evaluation as male donors, were selected by breeders at KWS based on their favorable yet contrasting cross-pollination performance on female plants in hybrid grain production trials. Selection was based on cross-pollination efficiency, assessed as grains per m^2^, g/m^2^, VAEX scores (0-9; where 0 indicates no VAEX and 9 indicates maximum VAEX). Further details on these traits are provided below and in Suppl. Table 1. Experimental hybrids were generated using the chemical hybridization agent Croisor® 100 (active ingredient sintofen; 1-(4-chlorophenyl)-5-(2-methoxyethoxy)-4-oxo-1,4-dihydrocinnoline-3-carboxylic acid; Asur Plant Breeding), applied to female plots to induce male sterility. All crossing blocks were conducted close to KWS Momont breeding station (48°17’36.3”N 1°36’53.4”E) in Allonnes, France in the seasons 2020-2021 and 2021-2022. At harvest, dry grain yield per plot was evaluated, thousand grain weight analysed and hybrid grain set expressed in number of grains per m^2^ calculated. Together with the hybrid grain-set, on the pollinator plots, maximum VAEX was scored, heading date (HD) in days after January 1^st^ was noted and plant height in cm (PH) was measured around BBCH 75. The 24 selected male lines from above were characterized by increased plant height and pronounced anther extrusion associated with high pollen production, whereas corresponding female lines exhibited reduced height, wider floral opening, and prolonged stigma receptivity to facilitate effective cross-pollination.

All 24 male-tester lines were also evaluated for their floral features under greenhouse and field conditions across two growth seasons (2021–2022 and 2022–2023) at the Leibniz Institute of Plant Genetics and Crop Plant Research (IPK), Gatersleben, Germany. Grains were germinated in soil trays at 20 °C and seedlings were vernalized for 8 weeks at 4 °C under a 12 h light/12 h dark photoperiod prior to transplantation into experimental environments.

In the greenhouse experiment, vernalized seedlings were transplanted into 13 × 13 cm pots and grown under long-day conditions (16 h light/8 h dark) at 18°C /15 °C Day/night until flowering. Each genotype was represented by two replications comprising 20 plants per replication. Field experiments were carried out at IPK, Gatersleben (51°49′39.0″ N, 11°16′30.1″ E) during the spring seasons of 2022 and 2023 (March-June). Seedlings were transplanted in silty loam soil plots, each consisting of five rows per genotype with a plant-to-plant spacing of 10 cm and row spacing of 15 cm. Standard agronomic and management practices were applied throughout the crop growth cycle, including irrigation twice weekly. In both environments, genotypes were evaluated in a completely randomized design with two replications.

### Phenotyping of male reproductive traits and floral bract features

To quantify variation in cross-pollination performance underlying hybrid grain set, traits related to anther extrusion dynamics, pollen release, and bract morphology were phenotyped during anthesis (Suppl. Table 1). Measurements captured complementary functional components of these reproductive processes using standardized imaging and sampling procedures. All traits were phenotyped using a Zeiss Stemi 508 stereomicroscope (Carl Zeiss Microscopy GmbH, Germany), except lemma–palea weight, which was determined gravimetrically. Main culm spikes were harvested at the onset of anthesis (W9.5), defined as the stage when the first anther emerged from florets of the central spikelets (S11–S13). Unless otherwise specified, measurements were conducted on second florets (F2) of the central spikelets following manual floret opening where required.

For each spike, three florets (one per spikelet) were sampled, and all three anthers per floret were analyzed when applicable. Three primary spikes were evaluated per replication, resulting in six spikes per genotype across two replications. All measurements were performed on independent biological replicates.

### Anther length (AL)

Anther length was measured at developmental stage Waddington stage 9.5 (Waddington *et al*., 1983) after removal of the lemma. Isolated anthers were imaged using a stereomicroscope equipped with ZEN Blue Lite software (version 3.3; Carl Zeiss Microscopy GmbH, Germany), and lengths were determined from captured images (Figure 1A).

**Figure 1.**
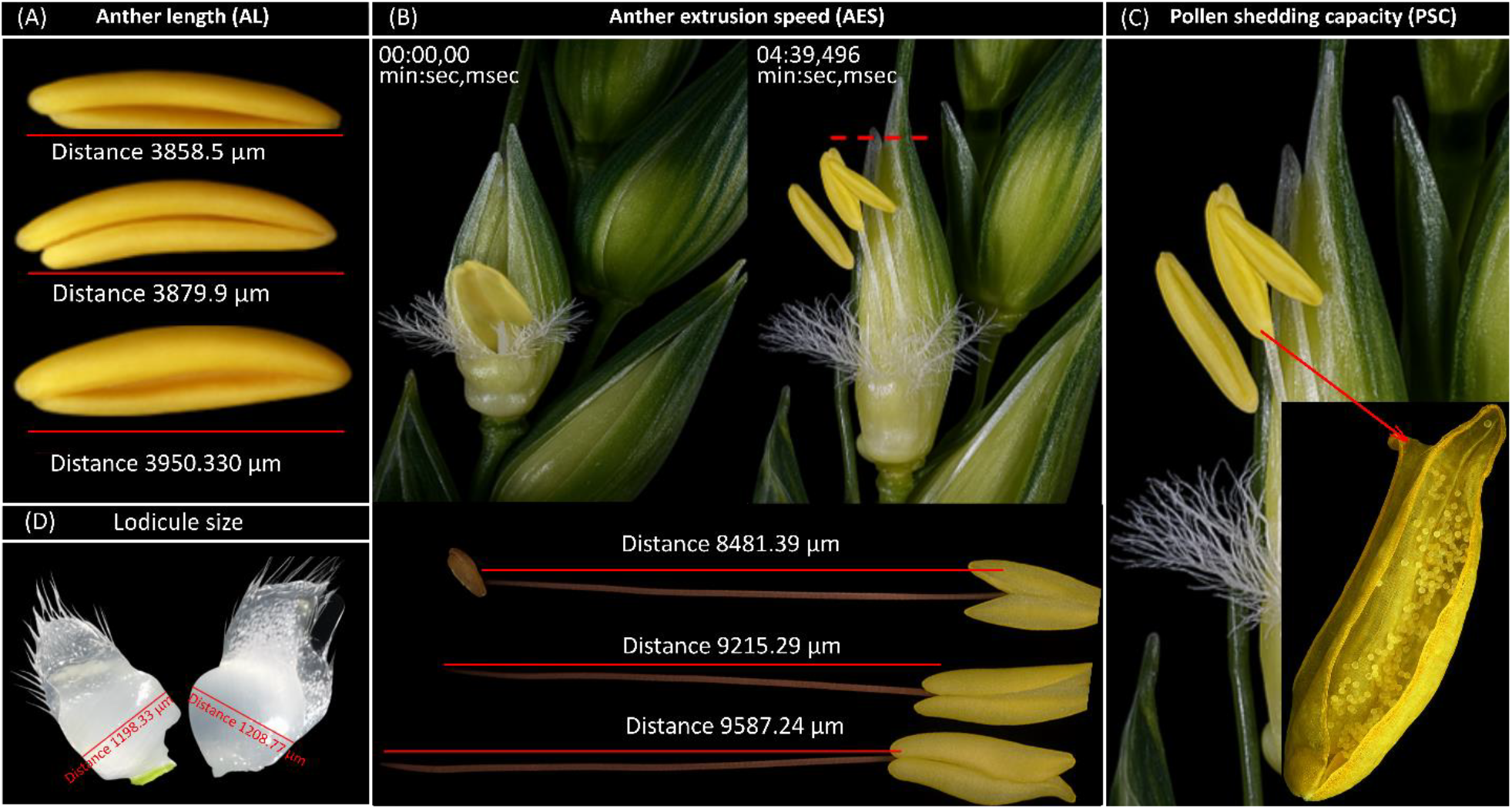
Representative images used for phenotyping of (A) anther length, (B) anther extrusion speed (red dashed line shows the extrusion stage), (C) pollen shedding, and (D) lodicule size. Distances are indicated in µm.

### Anther extrusion speed (AES)

Anther extrusion speed was calculated as a derived trait representing filament elongation per unit time. Florets were manually opened and positioned under a stereomicroscope to allow filament elongation during time-lapse imaging. The time required for anthers to reach the extrusion stage— defined as the point at which the anther tip reached the floret opening—was recorded. Anthers with filaments were then excised at the stamen–carpel junction, filament length was measured, and extrusion speed was calculated as filament length divided by elongation time (Figure 1B).

### Pollen shedding capacity (PSC)

Pollen shedding capacity was defined as the quantity of pollen available for cross-pollination per floret following anther dehiscence. At stage W9.5, florets were manually opened to expose mature anthers and permit filament elongation. Upon extrusion, anthers together with released pollen were collected and immediately transferred into Ampha 11 buffer. Pollen number was quantified using Ampha Z32 impedance flow cytometry (IFC, Amphasys AG, Switzerland) (Figure 1C).

### Lodicule width (LW)

Lodicule width was measured at developmental stage W9.5, when lodicules were fully swollen. Lodicules were carefully excised and imaged at high resolution, and their length was subsequently determined from the captured images (Figure 1D).

### Lemma palea weight (LPW)

Lemma (outer bract) and palea (inner bract) protect the floret’s reproductive organs. Lemma and palea weight were used as a proxy for lemma–palea strength, stiffness, and sturdiness. Five lemmas and five palea were collected from second florets of spikelets S10–S15 at stage W9.5. The collected tissues were pooled and weighed to determine combined lemma and palea weight. Three main culm spikes per genotype were analyzed per replication.

### Statistical analysis

Best linear unbiased predictors (BLUPs) for hybrid grain set and pollinator traits namely VAEX, plant height (PH), and heading date (HD), were estimated using linear mixed models fitted by restricted maximum likelihood (REML) (Patterson and Thompson, 1971; Smith *et al*., 2005). For hybrid grain set measured under field conditions, genotype, female parent, and year were included as random effects, whereas for pollinator traits, genotype and year were included as random effects. The respective models were:

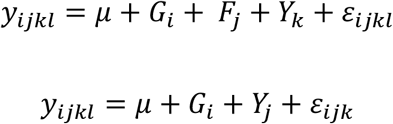

where *y* represents the observed trait value, *μ* the overall mean, *G* the genotype effect, *F* the female-parent effect, *Y* the year effect, and *ε* the residual error. Genotypic BLUPs were extracted from the fitted models for subsequent analyses.

For all the phenotypic traits, best linear unbiased estimators (BLUEs) were calculated across years and environments using a linear mixed model fitted REML. Genotype was treated as a fixed effect, whereas year, environment, replication nested within year × environment, and their interactions were fitted as random effects. The model applied was:

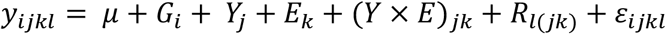

where y_ijkl_ denotes the observed trait value, μ the overall mean, G_i_ the genotype effect, Y_j_ the year effect, E_k_ the environment effect, (Y×E)_jk_ denotes the year × environment interaction, R_l(jk)_ the replication effect, and ε_ijkl_ the residual error. Linear mixed models were fitted using the *lme4* package in R, and BLUEs for genotypic effects were extracted from the fitted model.

Broad-sense heritability (*H*^*2*^) on a genotype mean basis was estimated from variance components obtained from a REML-based linear mixed model in which genotype, year, environment, and their interactions were treated as random effects. Heritability was calculated as:

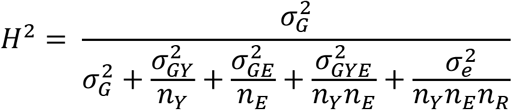

where *σ*^*2*^_*G*_ is genotypic variance, *σ*^*2*^_*GY*_, *σ*^*2*^_*GE*_, and *σ*^*2*^_*GYE*_ represent genotype interaction variances with year and environment, and *σ*^*2*^_*e*_ is the residual variance; *n*_*Y*_, *n*_*E*_, and *n*_*R*_ correspond to the numbers of years, environments, and replications, respectively (Falconer, 1996).

Trait relationships were first evaluated using Pearson’s correlation coefficients calculated from genotype BLUEs. To identify traits independently associated with hybrid grain set, multiple linear regression analysis was performed using BLUEs, with hybrid grain set fitted as the response variable and floral traits as predictors. Model selection was conducted using stepwise regression based on the Akaike information criterion (AIC) to identify the most parsimonious model (Akaike, 2003; Burnham and Anderson, 2002; Montgomery *et al*., 2021). Regression assumptions were assessed through residual diagnostics, and statistical significance was determined at *P* < 0.05.

Multicollinearity was evaluated before fitting multivariable regression models. Multicollinearity occurs when two or more predictor variables in the same model are strongly correlated and therefore describe partly overlapping variation, making it difficult to separate their individual effects on the response variable (Dormann *et al*., 2013). This was especially important in the present study because several male reproductive and floral traits were anticipated to be biologically interrelated. Multicollinearity was assessed using variance inflation factors (VIF) calculated with the *car* package in R; VIF values below 5 were considered indicative of acceptable collinearity (O’brien, 2007) (Suppl. Table 2). Standardized regression coefficients (β) were subsequently derived from the fitted regression model after standardizing variables to zero mean and unit variance, allowing comparison of relative trait contributions independent of measurement scale (Suppl. Table 3).

Multivariate phenotypic structure among genotypes was explored using principal component analysis (PCA) performed on standardized genotype BLUEs (mean = 0, SD = 1). Principal components were interpreted based on trait loadings.

Finally, path analysis was conducted using standardized regression coefficients to partition correlations between floral traits and hybrid grain set into direct and indirect effects, thereby quantifying coordinated trait influences on cross-pollination performance. All statistical analyses were performed in R.

## Results

### Genotypic and heritable trait variation underlies hybrid grain set

Considerable phenotypic variation was observed among the 24 elite winter wheat male tester-lines for floral and male reproductive traits. After cross-pollination, hybrid grain set on female plants (measured as grains m^-1^) ranged from 5,629 to 10,411 (mean = 7,811 ± 1576), indicating that cross-pollination efficiency varies greatly among male tester lines. We also found substantial variation in key male reproductive traits, including pollen shedding capacity (PSC, measured as pollen grains floret^-1^; 1,173-3,229; mean = 1,930± 478) and anther extrusion speed (AES, 803-6,295 µm min^−1^; mean = 2,132 ± 1214). Anther length (AL) varied between 3,103 and 5,472 µm (mean = 4,196 ± 496), whereas floral traits such as lodicule width (LW) as well as lemma and palea weight (LPW) exhibited comparatively narrower ranges, i.e. 972-1361; mean = 1168 ± 113 µm and 0.054-0.084; mean = 0.067 ± 0.01 g, respectively, indicating trait-specific variability among genotypes.

Broad-sense heritability (*H*^*2*^) estimates revealed strong genetic control of hybrid grain set, VAEX, and HD with *H*^*2*^ values of 0.88, 0.88, and 0.98, respectively. Likewise, reproductive traits showed high heritabilities, with AL exhibiting the highest heritability (*H*^*2*^ = 0.97), followed by AES (*H*^*2*^ *=* 0.89) and PSC (*H*^*2*^ *=* 0.88). LW (*H*^*2*^ *=* 0.83) and LPW (*H*^*2*^ *=* 0.76) displayed slightly lower but still substantial heritabilities. Overall, these results indicate that genetic factors accounted for a large proportion of the phenotypic variation in most traits across the evaluated environments.

Correlation analysis revealed strong associations among male reproductive traits (PSC, AES) associated with anther extrusion (VAEX) and hybrid grain set (Figure 2A). Hybrid grain set was positively correlated with the extent of VAEX (*r* = 0.70^***^), PSC; (*r* = 0.50^**^), and AES (*r* = 0.43^*^), indicating that both the magnitude of anther emergence and extrusion speed contribute to cross-pollination efficiency and hybrid grain set. AES, representing the speed at which anthers emerge during anthesis, showed positive correlations with anther length (AL; *r* = 0.65^***^) and PSC (*r* = 0.60^**^), suggesting that longer anthers reach floret tips slightly earlier and subsequently facilitate faster pollen shed. In contrast, VAEX, reflecting the degree of extruded anthers outside the floret, was strongly associated with AES (*r* = 0.71^***^) but exhibited an even stronger negative relationship with LPW(*r* = −0.75^***^; Figure 2A). This relationship indicates that while anther length influences extrusion dynamics, the final degree of anther extrusion is even more strongly constrained by bract features. For example, heavier lemma–palea tissues likely impose greater mechanical resistance during anthesis and floral opening, thereby limiting or hampering anther extrusion. Lower lemma–palea weight, however, may facilitate wider floret opening and thus allows a greater proportion of anthers to emerge. Consistent with such a floral bract constraint, we found that LPW was negatively correlated with AES (*r* = −0.55^**^) and hybrid grain set (*r* = −0.60^**^), supporting a trade-off between investment in protective floral tissues and cross-pollination efficiency. LW showed weak correlations with most of the traits (|*r*| ≤ 0.21; Figure 2A), indicating a comparatively minor independent contribution relative to the coordinated effects of extrusion dynamics and floral architecture.

**Figure 2.**
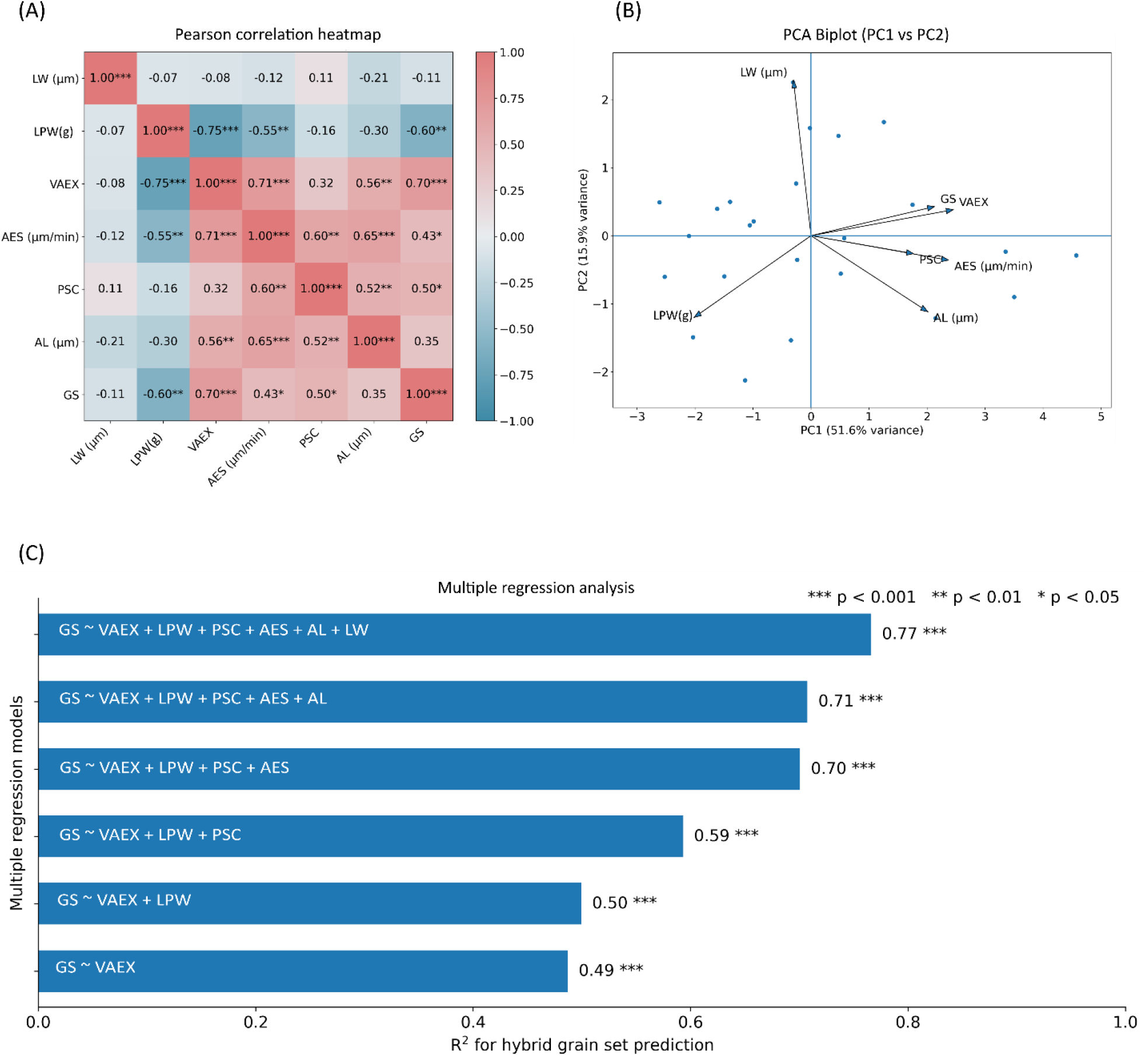
Multivariate analyses of floral traits associated with variation in hybrid grain set across wheat genotypes measured as a proxy output trait for cross-pollination efficiency (A) Heatmap of Pearson correlation coefficients among floral traits and hybrid grain set calculated from genotype BLUEs. Cell colors represent correlation magnitude and direction, and numerical values indicate Pearson’s correlation coefficients (r). (B) Principal component analysis (PCA) biplot illustrating multivariate relationships among floral traits and genotypes. PC1 and PC2 explain 51.6% and 15.9% of total variance, respectively. Arrows represent trait loadings and points indicate individual genotypes. (C) Sequential multiple regression models showing the increase in explained variance (R^2^) for hybrid grain set prediction as additional traits were incorporated. Bars represent model R^2^ values.; statistical significance is denoted by asterisks (p < 0.05, p < 0.01, p < 0.001). Abbreviations: GS - hybrid grain set; LW - lodicule width; LPW - lemma palea weight, VAEX-visual anther extrusion; AES-anther extrusion speed; PSC-pollen shedding capacity; AL-anther length.

### Key male reproductive traits independently determine the efficiency of hybrid grain production

Multiple regression analysis demonstrated that floral traits collectively explained a large proportion of variation in hybrid grain set among genotypes (*R*^*2*^ = 0.763, *P* < 0.001), indicating strong multivariate regulation of cross-pollination performance. Among predictors, VAEX and PSC exhibited significant positive effects on hybrid grain set, with VAEX showing the largest contribution with highest standardized beta (std. *β* = 0.73, *t* = 3.25, *P* = 0.001; Suppl. Table 3) and PSC also contributing significantly (std. *β* =0.71, *t* = 4.27, *P* = 0.01; Suppl. Table 3). The concurrent significance of these traits indicates that successful hybrid grain set is primarily associated with successful anther extrusion (VAEX) coupled with pollen release outside the floret (PSC), linking extrusion efficiency directly to cross-pollination and fertilization success. In contrast, AES displayed a significant negative standardized effect (std. *β* = −0.61, *t* = −2.81, *P* = 0.01; Suppl. Table 3) despite its positive pairwise association with hybrid grain set, indicating shared variance among extrusion-related traits. This result suggests that AES functions primarily as an upstream kinetic component for other traits to influence hybrid grain set indirectly. LPW, LW, and AL, however, displayed weaker negative standardized betas (Suppl. Table 3), suggesting that these traits contributed less independently to variation in grain set once PSC and VAEX were included in the model.

Sequential model construction revealed a progressive increase in explanatory power as male reproductive traits were incorporated, with model fit improving markedly following the inclusion of VAEX and PSC (Figure 2C), highlighting their central contributions to the observed variation for hybrid grain set.

Variance inflation factor (VIF) analysis confirmed our model stability with all predictors exhibiting values below 5 (Suppl. Table 2). Moderate collinearity observed for VAEX (VIF = 3.70) and AES (VIF = 3.38) likely reflects biological integration among extrusion-related traits rather than statistical redundancy. Collectively, these results indicate that the observed variation in hybrid grain set is structured by a hierarchical interaction among floral traits: while extrusion dynamics and pollen shedding represent the primary functional determinants for variation in hybrid grain set, structural attributes such as lemma–palea weight impose secondary floral constraints on cross-pollination efficiency.

### Multivariate trait analysis distinguishes cross-pollination performance from floral structural investments

Principal component analysis (PCA) captured major patterns of multivariate phenotypic variation among genotypes, with the first two components explaining 67.5% of total variance (PC1: 51.6%; PC2: 15.9%; Figure 2B). PC1 was defined primarily by coordinated variation among male reproductive traits, including VAEX, PSC, AES, AL, and cross-pollination performance measured as hybrid grain set. All of these traits loaded positively along the PC1 axis, whereas LPW loaded in the opposite negative direction, indicating contrasting contributions of male reproductive functions and floral structural investments. Accordingly, PC1 differentiated genotypes along a gradient of cross-pollination performance (Figure 2B). PC2 was driven mainly by variation in lodicule width, which was oriented largely independently from other traits, indicating comparatively distinct phenotypic variation for this trait (Figure 2B). The PCA biplot therefore illustrates that multivariate structuring among genotypes is governed positively predominantly by coordinated male reproductive traits (cross-pollination ability) together with negative structural floral attributes LPW.

### Direct and indirect trait interactions shape cross-pollination ability and hybrid grain set

Path analysis partitioned trait associations into direct and indirect components, clarifying the functional pathways underlying variation in cross-pollination ability and hybrid grain set (Figure 3). VAEX exhibited the strongest positive direct effect (0.73;) with minimal indirect influence (-0.02), resulting in the largest overall contribution (total effect = 0.70; Suppl. Table 4), consistent with its primary role as the immediate determinant of cross-pollination success. PSC also showed a substantial positive direct effect (0.70), partially moderated by negative indirect effects (−0.21), yielding a strong net contribution (total effect = 0.50; Suppl. Table 4) and indicating that better PSC directly promotes pollen dispersal and cross-pollination. In contrast, AL did not exert a positive independent effect on hybrid grain set, despite showing a positive overall association with the trait. Path analysis revealed a small negative direct effect (direct effect = -0.19; Suppl. Table 4), but a stronger positive indirect effect through other reproductive traits (indirect effect = 0.54; Suppl. Table 4). This suggests that the apparent contribution of AL to hybrid grain set is mediated largely through its association with PSC, VAEX, and AES, indicating that AL may support grain set indirectly as part of a coordinated reproductive phenotype rather than acting as a primary direct driver of cross-pollination success. Similarly, AES displayed a pronounced negative direct effect (−0.61) but a large positive indirect effect (1.04), producing an overall positive association (total effect = 0.44; Suppl. Table 4). These trait effects indicate that the contribution of AES is mediated primarily through its upstream effects on the timing of VAEX and on PSC, rather than through an independent direct pathway.

**Figure 3.**
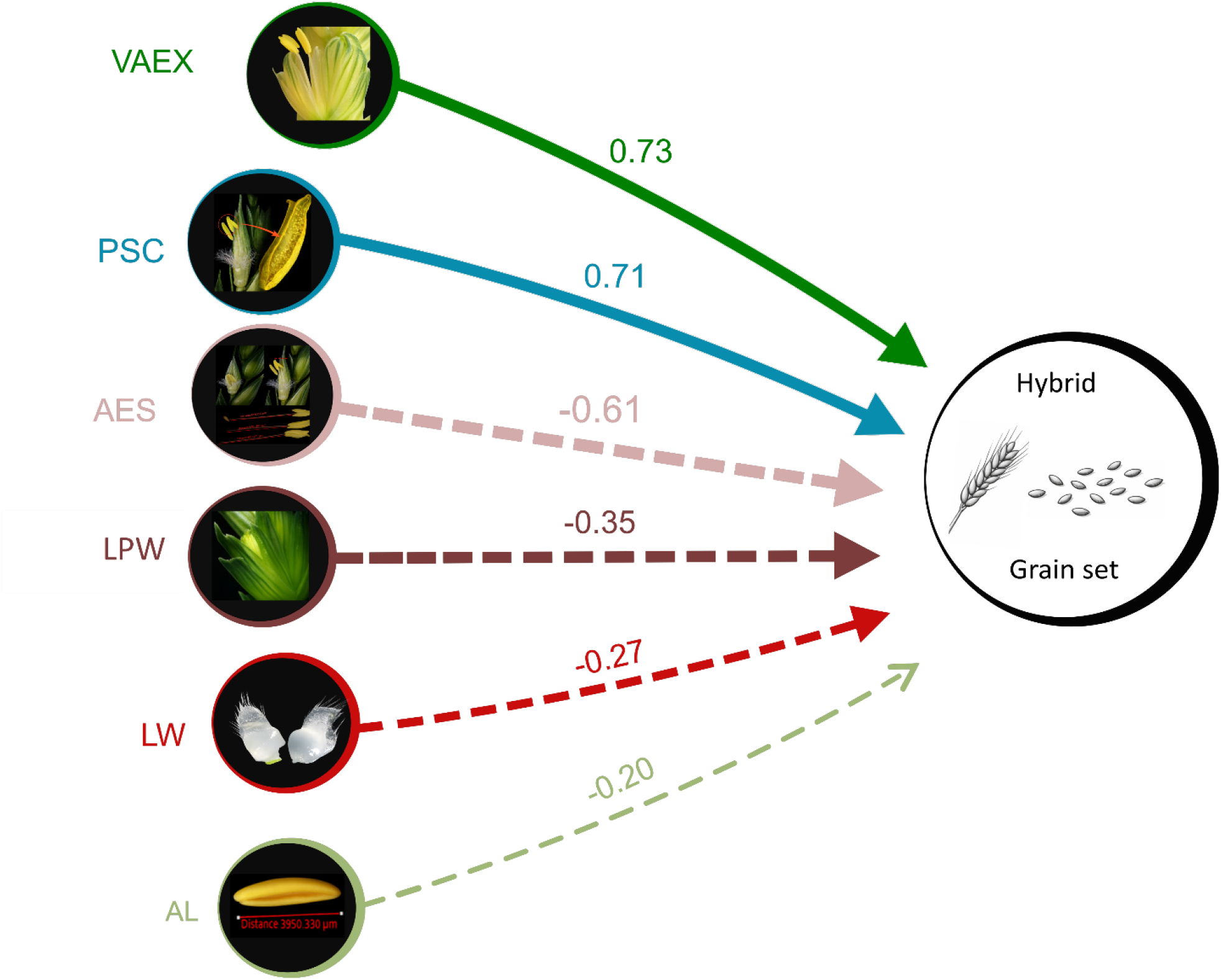
Path diagram illustrating the direct effects of floral traits on reproductive performance in wheat based on standardized path coefficients. Arrows represent direct relationships between traits and hybrid grain set, with arrow thickness proportional to effect magnitude. Solid arrows indicate positive effects, whereas dashed arrows indicate negative effects. Values shown on arrows correspond to standardized direct effect estimates derived from path analysis.

Morphological bract traits contributed predominantly negative effects. LPW exerted both negative direct (−0.35) and indirect (−0.24) influences, resulting in the strongest overall negative contribution (total effect = −0.60; Suppl. Table 4). This pattern supports a trade-off between structural bract investment enclosing floral tissues and cross-pollination function, whereby increased floral bract weight likely imposes mechanical resistance to floret opening and constrains effective anther extrusion. In contrast, lodicule width showed comparatively weak effects (direct = −0.26; indirect = 0.15; total effect = −0.11; Suppl. Table 4), indicating limited independent involvement in determining hybrid grain set outcomes. Collectively, these results demonstrate hybrid grain set as a proxy output of cross-pollination ability and is governed primarily by direct effects of traits associated with pollen release and anther extrusion, whereas structural bract attributes and extrusion kinetics influence cross-pollination outcomes indirectly through coordinated trait interactions.

## Discussion

### Hybrid grain production is determined by cross-pollination components rather than extrusion magnitude alone

Studies have repeatedly identified that VAEX is a fundamental precondition and key floral trait influencing hybrid grain production in wheat. Quantitative genetic analyses have shown that variation in anther extrusion is heritable and associated with hybrid grain set (Boeven *et al*., 2016; El Hanafi *et al*., 2022). However, these studies also indicate that the relationship between VAEX and cross-pollination performance is incomplete, suggesting that additional reproductive factors contribute to hybrid grain set (Adhikari *et al*., 2020; Winn *et al*., 2021). Our results are consistent with this interpretation. While VAEX showed a positive association with hybrid grain set, incorporating additional traits substantially increased the proportion of explained variation, indicating that extrusion alone captures only part of the cross-pollination processes underlying hybrid grain production. This motivates dissection of the processes in addition to anther extrusion that determine effective pollen export.

Among all traits examined, PSC exerted the strongest direct positive effect on hybrid grain set, indicating that effective pollen release is a more immediate determinant of fertilization success than extrusion magnitude alone. In predominantly autogamous and often cleistogamous wheat florets, anther dehiscence initiates during filament elongation, before complete extrusion (Heslop-Harrison, 1968; Browne et al., 2018; Selva et al., 2020). Consequently, a proportion of pollen is discharged within the floret prior to external exposure. The amount of pollen available for cross-pollination after extrusion therefore depends on both the timing of dehiscence and the ability of anthers to overcome the mechanical resistance imposed by the enclosing bracts—lemma and palea (Whitford *et al*., 2013; Zajączkowska *et al*., 2021). Genotypic differences in these processes likely determine the quantity of pollen effectively exported, which may explain why genotypes with similar levels of anther extrusion can nevertheless differ markedly in hybrid grain production. This is illustrated by KWS_04 and KWS_09, which share the same VAEX ranking 6^th^ but differ considerably in hybrid grain set ability, ranking 15^th^ and 9^th^, respectively (Suppl. Table 1). Within this context, PSC captures the functional component of pollen dispersal that directly contributes to hybrid grain set.

The high heritability observed for PSC highlights its potential as a robust selection target for hybrid wheat breeding. Previous studies have reported substantial genetic variation in male floral traits associated with pollen production and dispersal (El Hanafi *et al*., 2022; Garst *et al*., 2023), indicating that components of pollen export are amenable to genetic improvement. Our results extend these findings by identifying pollen shedding capacity as a key functional trait linking floral morphology to cross-pollination success and hybrid grain set.

### Extrusion kinetics affect hybrid grain set indirectly

AES showed a positive association with hybrid grain set at the correlation level but a negative direct effect in the regression and path analyses. This apparent contradiction reflects shared variance among extrusion-related traits and suggests that AES functions primarily as an upstream kinetic component to VAEX and PSC rather than as an independent determinant of reproductive success. Biologically, rapid extrusion may facilitate timely pollen presentation, yet without sufficient pollen shedding capacity, or low enclosure resistance, or an adequate extent of anther extrusion, increased speed alone does not ensure effective pollen dispersal. Previous studies have shown that successful pollen dispersal in wheat depends on coordinated timing between anther emergence, dehiscence, and pollen release (Selva *et al*., 2020; Zajączkowska *et al*., 2021). The large positive indirect effect of AES suggests that extrusion kinetics promote cross-pollination performance only when integrated with downstream processes like VAEX and PSC governing pollen release and exposure.

### Floral bract investment constrains reproductive performance

Lemma–palea weight exhibited a consistent negative association with extrusion traits and hybrid grain set, supporting a mechanical constraint hypothesis. Heavier floral bracts likely impose greater resistance to floret opening and anther emergence, thereby limiting effective pollen exposure. Similar structural constraints have been described in wheat floral biology, where the extent of floret opening is determined by the interaction between lodicule expansion and floral bracts (Browne *et al*., 2018; Selva *et al*., 2020; Szeliga *et al*., 2023).

Such structural trade-offs provide a plausible explanation for why some genotypes with an adequate extent of anther extrusion fail to achieve high hybrid grain set. Even when extrusion speed is favorable, incomplete floret opening may restrict pollen dispersal e.g. KWS_04 (higher LPW; lower PSC; poor GS) and KWS_09 (low LPW; higher PSC; good GS) (Suppl. Table 1). These findings highlight the importance of considering floral bract architecture when evaluating male parental lines and suggest that structural traits should be integrated alongside reproductive dynamics in phenotyping pipelines (Selva *et al*., 2020; Whitford *et al*., 2013)

### Multivariate trait architecture underlies hybrid grain production

Our principal component and regression analyses indicate that hybrid grain set is primarily structured by coordinated male reproductive traits, whereas floral bract traits contribute largely independent variation. This pattern suggests that male reproductive performance and floral bract architecture represent distinct components of phenotypic variation during anthesis. Consistent with this interpretation, the multivariate framework explained substantially more variation in hybrid grain set than single-trait evaluations. Previous studies have similarly shown that male fertility traits in wheat are controlled by complex quantitative architectures involving multiple interacting loci and phenotypic components (Boeven *et al*., 2016; Singh *et al*., 2021). Integrating these results suggests a hierarchical organization of male reproductive processes in which floral structures influence floret opening, extrusion dynamics, determine anther exposure, pollen shedding capacity, and ultimately regulate the amount of pollen released for dispersal. Within this framework, male reproductive performance emerges from interacting floral processes rather than from any single trait.

### Implications for hybrid wheat breeding

From a breeding perspective, these findings highlight the need to extend male line phenotyping beyond single-trait evaluation of anther extrusion. Although VAEX remains a useful initial screening trait, its predictive value increases when considered together with pollen shedding capacity and floral structural characteristics. The high heritability observed for several male reproductive traits further indicates that components of pollen export efficiency are amenable to genetic improvement. As hybrid wheat breeding increasingly focuses on optimizing floral biology to enhance cross-pollination efficiency (Rohde *et al*., 2025; Selva *et al*., 2020; Whitford *et al*., 2013), integrating multivariate trait assessment into breeding pipelines could facilitate more reliable identification of superior male parental lines and improve hybrid grain production efficiency.

### Conclusions and current limitations

This study was conducted within a defined germplasm panel and environmental context, and the stability of the observed trait relationships remains to be tested across wider genetic backgrounds and environments. Further investigation of the genetic basis of pollen shedding capacity and anther extrusion dynamics may facilitate the development of marker-assisted or genomic selection strategies for reproductive efficiency (Boeven *et al*., 2016; Singh *et al*., 2021). Because many of these traits are microscopic and therefore difficult to phenotype rapidly at scale, the identification of reliable proxy traits, including anther length and lemma–palea weight, together with the development of higher-throughput phenotyping approaches, will be important. Their evaluation in larger hybrid grain production trials will further clarify their utility under commercial breeding conditions.

## Supplementary data

All the supplementary data is being provided as a separate excel file.

## Acknowledgement

We thank Dr. Antje Rohde (KWS Einbeck, Germany) for critically reading previous versions of the manuscript as well as Kerstin Wolf and Corinna Trautewig for their excellent technical assistance. We also acknowledge the members of the Cryo and Stress Biology group for their support during phenotyping and IFC measurements. Finally, we extend our gratitude to Peter Schreiber and the greenhouse and field management team for their dedicated care of the plants throughout the study and for maintaining the experimental facilities at IPK, including IPK infrastructure. While conducting this study TS received financial support from the Federal Ministry of Education and Research (BMBF).

## Author contribution

DK: Conceptualization, Methodology, Investigation, Data curation, Formal analysis, Data interpretation, Validation, Visualization, Writing – original draft, Writing – review & editing; MS: Resources, Data interpretation, Writing – review & editing; LG: Resources, Data curation, Writing – review & editing; TS: Conceptualization, Funding acquisition, Project administration, Supervision, Validation, Writing – review & editing.

All authors reviewed the manuscript, contributed to revisions, and approved the final version

## Data availability

All the data, tables and figures generated are original and available within the paper, supplementary files and upon request from the corresponding authors.

